# Pre-existing YFV-17D immunity mediates T cell cross-protection against DENV-2 infection

**DOI:** 10.1101/2025.04.16.649113

**Authors:** Prince Baffour Tonto, Sebastian Gallon, Reem Alatrash, Wei-Kung Wang, Bobby Brooke Herrera

**Affiliations:** Rutgers Global Health Institute, Rutgers University, New Brunswick, NJ, USA; Department of Medicine, Division of Allergy, Immunology, and Infectious Diseases and Child Health Institute of New Jersey, Rutgers Robert Wood Johnson Medical School, New Brunswick, NJ, USA; Department of Tropical Medicine, Medical Microbiology and Pharmacology, John A. Burns School of Medicine, University of Hawaii at Manoa, Honolulu, HI, USA

## Abstract

Widespread yellow fever virus (YFV) immunity in Sub-Saharan Africa may mitigate orthoflavivirus outbreaks. Here, we investigate whether pre-existing YFV-17D immunity confers cross-protection against dengue virus serotype 2 (DENV-2) in a murine model. *IFNAR1^-/-^* mice immunized with YFV-17D exhibited significantly reduced DENV-2 viremia, weight loss, and disease severity, with improved survival compared to YFV-naïve controls. Mechanistic studies revealed that cross-protection was mediated by heterologous T cell responses rather than cross-neutralizing antibodies. Depletion of T cells in YFV-17D-immune mice prior to DENV-2 challenge resulted in increased viremia, weight loss, and disease severity, underscoring the protective role of YFV-17D-elicited T cell immunity. Furthermore, YFV-17D-specific T cells displayed cytotoxicity against DENV NS3- and NS5-pulsed cells, demonstrating their functional role in viral control. These findings highlight the critical contribution of heterologous T cell immunity in YFV-17D-mediated protection against DENV-2 and suggest that vaccines designed to elicit T cell responses could enhance cross-protection against orthoflavivirus infections.

## Introduction

Orthoflaviviruses present a significant global health challenge due to their substantial annual disease burden (1). Despite the availability of vaccines for certain orthoflaviviruses, such as yellow fever virus (YFV), dengue virus serotypes 1-4 (DENV-1-4), and Japanese encephalitis virus (JEV), recent studies suggest immunization strategies may be complicated by heterologous immune interactions, which can either confer cross-protection or enhance disease severity in endemic regions (1–6). While the live-attenuated yellow fever virus 17D (YFV-17D) vaccine has provided significant protection against yellow fever for over eight decades, its impact on immunity to other orthoflaviviruses remains underexplored (7–9).

A defining feature of orthoflaviviruses is their high degree of antigenic similarity, which facilitates heterologous immune responses between related viruses (10). This cross-reactivity has been extensively studied in the context of DENV and Zika virus (ZIKV), where prior exposure to one virus can shape the immune response to subsequent infection with another (11–22). While heterologous DENV or ZIKV immunity can provide cross-protection(12, 13, 18, 19, 21, 23–25), it can also exacerbate disease severity via antibody-dependent enhancement (11, 20, 26). Given the broad epidemiological footprint of orthoflaviviruses, their epidemic and pandemic potential, and the protracted timelines required for vaccine development and regulatory approval, understanding how pre-existing immunity influences subsequent heterotypic infections is essential (25). Moreover, this immunological cross-reactivity may provide insights into epidemiological patterns, including the apparent absence or limited transmission of specific orthoflaviviruses in regions where related viruses are endemic (23, 24). Additionally, it may explain the predominance of mild or clinically inapparent outbreaks in areas where multiple orthoflaviviruses co-circulate, as pre-existing immunity could modulate disease severity (27).

Asia, the Americas, and Africa bear the highest burden of orthoflavivirus-related diseases (1, 28–30). However, a long-standing epidemiological paradox is the lack of urban YFV transmission in South America and the absence of YFV in Asia, despite the presence of an immunologically naïve population and a high density of *Aedes aegypti*, the primary mosquito vector for YFV (31). Given the hyperendemicity of DENV and ZIKV in these regions, it has been hypothesized that heterologous immunity from prior DENV and ZIKV infections might confer cross-protection against YFV. This hypothesis is supported by preclinical studies demonstrating that mice and macaques with pre-existing immunity to DENV and ZIKV exhibit reduced YFV viremia compared to naïve controls (23, 24).

Conversely, in Sub-Saharan Africa, orthoflaviviruses such as YFV, DENV, ZIKV and West Nile virus (WNV) co-circulate endemically (32–34). Interestingly, despite the co-circulation of these viruses, outbreaks in this region tend to be mild and clinically inapparent (27, 30). One possible explanation is the widespread pre-existing immunity to YFV, primarily driven by extensive vaccination campaigns and, to a lesser extent, natural infections. This pre-existing YFV immunity may modulate the severity and clinical outcomes of subsequent heterotypic orthoflavivirus infections. However, the precise immunological mechanisms underlying this phenomenon remain incompletely understood and warrant further investigation.

In this study, we hypothesized that pre-existing YFV-17D immunity confers cross-protection against subsequent DENV-2 infection. Using a murine model, we aimed to evaluate the impact of YFV-17D immunity on DENV-2 disease severity, clinical outcomes, and the underlying immune mechanisms driving YFV-17D-mediated cross-protection.

## Methods

### Animal care, ethics statement and in vivo study design

All experimental procedures involving animals were conducted in accordance with institutional guidance and approved by the Rutgers, The State University of New Jersey Institutional Animal Care and Use Committee (IACUC) (protocol approval number 202300007). Efforts were done to minimize animal suffering, including humane endpoints and appropriate monitoring. *IFNAR1^-/-^* mice were purchased from Jackson Laboratories and maintained in a breeding colony at the Child Health Institute of New Jersey, part of the Rutgers Robert Wood Johnson Medical School. Five-week-old mice were sham-immunized (saline) or immunized twice, one week apart, with 10^4^ PFU of YFV-17D via subcutaneous injection. Fourteen days following the second immunization, mice were either humanely euthanized using CO2 exposure and cardiac puncture for blood and spleen collection or challenged retro-orbitally with a lethal dose (2×10^7^ PFU) of the mouse adapted strain DENV-2 S221. To assess viremia, submandibular blood samples were collected on days 2 and 4 post-DENV-2 challenge. Mice were monitored daily for weight loss and clinical symptoms until day 6, while survival was recorded up to day 14 post-infection.

### Cells

Vero CCL-81 (Kidney epithelial cells isolated from an African green monkey, ATCC) cells were cultured in an Eagle’s Minimal Essential Medium with L-glutamine, supplemented with 10% heat-inactivated fetal bovine serum (FBS) and 1% penicillin-streptomycin solution. U937 (pro-monocytic cells isolated from a patient with histiocytic lymphoma, ATCC) cells were cultured in RPMI-1640 medium with L-glutamine supplemented with 10% heat-inactivated fetal bovine serum (FBS) and 1% penicillin-streptomycin solution. Both cell lines were maintained at 37°C in a humidified incubator with 5% CO2.

### Viruses and titer determination

All viruses, including YFV-17D, ZIKV (PRVABC59), and WNV (Bird 114) were obtained from BEI Resources, except DENV-2 S221, which was kindly provided by Dr. Sujan Shresta from La Jolla Institute for Immunology. All viruses were propagated in Vero cells and viral titers were determined using the TCID50 assay as previously described (35).

### RNA extraction and reverse transcriptase PCR

Viral RNA was extracted from plasma using the QIAamp^®^ Viral RNA Mini Kit, following the manufacturer’s instructions (QIAGEN). The extracted RNA was then reverse transcribed into cDNA and amplified in a one-step real-time polymerase chain reaction (PCR), with a master mix and primer-probe mixture from Integrated DNA Technology. The thermal cycling reactions conditions were as follows: reverse transcription, 50^0^C for 15 min; polymerase activation, 95^0^C for 3 min; denaturation, 95^0^C for 15 sec; annealing/extension, 60^0^C for 1 min. A total of 40 amplification cycles were performed. The primers and probe used for DENV-2 detection were obtained from previously published literature (36). To quantify viral loads, a DENV-2 standard curve was generated using genomic RNA from BEI Resources. The number of DENV-2 RNA copies was calculated using the genome equivalents formula: [ssRNA concentration (g/µL) * 6.022 × 10^23^(copy/mol)]/ [ssRNA length * 340 (g/mol)]. A stock solution of 10^8^ RNA copies/µL was prepared and serially diluted 10-fold to generate working solutions ranging from 10^0^ to 10^7^ copies/µL.

### Microneutralization assay

Microneutralization assays were performed to detect neutralizing antibodies against YFV-17D, DENV-2, ZIKV and WNV, as previously described (34, 37). Plasma was diluted in a two-fold series from 1:40 to 1:2560 in EMEM supplemented with 2% FBS and L-glutamine. The dilutions were prepared in a 96-well plate, with each well incubated with 100 plaque forming units (PFU) of the respective virus for one hour at 37°C in a 5% CO2 incubator. After incubation, the virus-serum complexes were then added to Vero CCL-81 cells (2 x 10^4^ cells/well) and further incubated for 5 days for YFV-17D, ZIKV and WNV, and 10 days for DENV-2. Cytopathic effects were assessed by staining with 0.2% crystal violet (SigmaAldrich) stain. Following 24 hours of staining, the plates were washed with copious amounts of tap water. For negative controls, at least 2 orthoflavivirus-naïve plasma samples per plate were incubated with the viruses prior to addition to Vero cells. For positive controls, 100 PFU of each virus was plated on Vero cells without prior serum contact, in duplicate per plate. The neutralizing titer was defined as the inverse of the highest serum dilution that resulted in a 90% reduction in infection. A titer <1:40 was considered negative, while a titer ≥1:40 was considered positive.

### Viral lysate Western blot

Viral lysates were prepared from virus-infected Vero cells as previously described (33, 38). Briefly, the lysates were subjected to SDS-PAGE to separate viral proteins based on molecular weight. The resolved proteins were then transferred onto PVDF membrane and probed with plasma from YFV-17D-immunized mice (1:300 dilution). Following an overnight serum incubation, the membrane was treated with goat anti-mouse IgG-HRP secondary antibody (1:2000). Viral protein detection was performed using the Clarity^TM^ Western ECL chemiluminescent substrate (BioRad), followed by image acquisition using the ChemiDoc^TM^ Imaging System (BioRad).

### Antibody dependent enhancement (ADE)

The ADE assay was performed as previously described (11). In brief, plasma was diluted in a 10-fold series from 1:10 to 1:100,000 in RPMI-1640 medium supplemented with 2% FBS. Each dilution was incubated with a multiplicity of infection (1 MOI) of YFV-17D, 5 MOI of DENV-2, 2 MOI of ZIKV, and 0.5 MOI of WNV for one hour at 37°C in a 5% CO2 incubator. Following incubation, the virus-plasma antibody complexes were then added to U937 cells (3 x 10^4^ cells/well), followed by a 2-hour incubation at 37°C with 5% CO2. Afterwards, the U937 cells were resuspended in RPMI-1640 supplemented with 10% FBS, followed by incubation at 37°C with 5% CO2 for varying durations depending on the virus: 2 days for YFV-17D, 4 days for DENV-2 and ZIKV, and 1 day for WNV. The cells were then fixed, permeabilized and stained intracellularly with 4G2-Alexa flour 532, followed by flow cytometry analysis, with the acquisition of 2×10^4^ events to identify population of infected cells as an indication of infection enhancement. The gating strategy is shown in Supplementary Fig 1.

### Evaluation of cross-reactive YFV 17D-specific T-cells

2×10^6^ splenocytes, prepared as a single cell suspension from spleen tissue, were pulsed with 10 μg/ml of a modified version of the *Bacillus anthracis* lethal factor (LFn) fused to structural (capsid) and non-structural proteins (NS3, NS4b, NS5-MTase, NS5-RdRp) (purchased from Mir Biosciences, Inc.) from both YFV and DENV, followed by incubation for 24 hours at 37°C with 5% CO2. The cells were then stained extracellularly with fluorophore-conjugated antibodies specific for viable cells, CD3^+^ T cells (Alexa flour 700-anti mouse CD3, Invitrogen), B cells (BV 650-anti mouse CD19, Invitrogen), CD4^+^ (APC-eflour 780-anti mouse CD4, Invitrogen) and CD8^+^ (Alexa flour 488-anti mouse CD8a, Invitrogen) T cells. Afterwards, the cells were fixed with fixation buffer (Invitrogen), permeabilized with permeabilization buffer (Invitrogen) and stained intracellularly with fluorophore-conjugated antibody specific for IFNγ-producing cells (eflour 660-anti mouse IFNγ, Invitrogen). Flow cytometry analysis was conducted with the acquisition of 5×10^5^ events using a Cytek Aurora instrument (Cytek Biosciences) to identify the population of T cell subsets. The gating strategy is shown in Supplementary Fig. 2.

### CD4^+^ and CD8^+^ T cell depletion

5-week-old mice were sham-immunized or immunized twice, one week apart with 10^4^ PFU of YFV-17D virus via subcutaneous injection. The mice were allowed to build immunity for 14 days following the second immunization. Prior to retro-orbital challenge with a lethal dose of DENV-2 S221 (2 × 10⁷ PFU), the mice received an intraperitoneal injection of 200 µg of either IgG2a isotype control (Bioxcell), anti-CD4 (anti-mouse CD4 clone: GK 1.5, Bioxcell) or anti-CD8α (anti-mouse CD8α clone: 2.43, Bioxcell) antibodies at 3 days and 1 day before the challenge, and 1 day post-DENV-2 infection. Mice were bled on days 2 and 4 following DENV-2 challenge to determine viremia. They were monitored for weight change, clinical symptoms, and survival until day 4 post DENV-2 infection, at which point they were euthanized for spleen harvest to confirm CD4^+^ and CD8^+^ T cell depletion. To confirm depletion, splenocytes were stained extracellularly with fluorophore-conjugated antibodies specific for viable cells, CD3^+^ T cells (Alexa flour 700-anti mouse CD3, Invitrogen), B cells (BV 650-anti mouse CD19, Invitrogen), CD4^+^ (APC-eflour 780-anti mouse CD4, Invitrogen) and CD8^+^ (Alexa flour 488-anti mouse CD8a, Invitrogen) T cells. Flow cytometry analysis was conducted with the acquisition of 5×10^5^ events using a Cytek Aurora instrument (Cytek Biosciences) to identify the population of T cell subsets. The gating strategy is shown in Supplementary Fig. 3.

### In vivo cytotoxicity assay

The in vivo cytotoxicity assay was performed as previously described (39). Briefly splenocytes from YFV-naïve mice were pulsed with either 10 μg/ml of LFn-conjugated NS3 and NS5 proteins of DENV or RdRp of La Crosse virus (LACV) (irrelevant immunogen) (purchased from Mir Biosciences, Inc.) for 24 hours. Cells were washed and then stained with Carboxyflourescein succinimidyl esther CFSE (Invitrogen) in 1X PBS for 10 min at 37°C. To distinguish between the targets, cells pulsed with DENV non-structural proteins were stained with of CFSE (CFSE^High^), while those pulsed with LACV-RdRp were stained with 0.5 of CFSE (CFSE^Low^). Following staining, CFSE^High^ and CFSE^Low^ cells were mixed in a 1:1 ratio, and adoptively transfer into YFV immunized or naïve mice via retro-orbital injection. Sixteen hours later, splenocytes were collected to quantify CFSE-labelled cells by flow cytometry, with the acquisition of 10^6^ events. The specific percentage killing was calculated as follows: 100 − ((% DENV NS3 or NS5-pulsed cells in YFV-immune mice / % LACV-RdRp pulsed cells in YFV-immune mice) / (% DENV NS3 or NS5-pulsed cells in naïve mice / % LACV-RdRp pulsed cells in naïve mice)) x 100.

### Clinical scoring of disease

Clinical scoring criteria included signs such as ruffled fur, hunched posture, lethargy and death. The basis of clinical features was done using a 5-point scale: 1, healthy; 2, ruffled fur; 3, ruffled fur and hunched posture; 4, ruffled fur, hunched posture and lethargy; 5, dead (Supplementary Table 1).

### Statistical analysis

The normality of the data was assessed using the Kolmogorov-Smirnov test (40). Differences between two dependent and independent groups were assessed using the two-tailed Wilcoxon signed-rank test or the Mann-Whitney test, respectively. For parametric data, comparisons between more than two independent groups were assessed using one-way ANOVA, followed by Tukey’s multiple comparison or Holm-Šídák’s multiple comparisons test. For non-parametric data, comparisons between more than two independent groups were assessed using the Kruskal-Wallis test followed by Dunn’s test for multiple comparisons. Differences in weight loss between two or more than 2 groups over time were analyzed using two-way ANOVA with a mixed-effect model for multiple comparisons. Cut-off values for identifying CD4^+^IFNγ^+^ and CD8^+^IFNγ^+^ positive responses were determined using receiver operating characteristic curve analysis (ROC) (41). All statistical analyses were performed using GraphPad Prism 10.

## Results

### Pre-existing YFV-17D immunity provides protection against DENV-2 infection

To evaluate the effect of pre-existing YFV-17D immunity on subsequent DENV-2 infection, *IFNAR1^-/-^* were either sham-immunized or immunized twice with YFV-17D at a 7 day interval. 14 days after the second immunization, mice were challenged with DENV-2 S221 (Fig. 1A). By 2 days post-infection (dpi), YFV-17D-naïve mice exhibited an average body weight loss of 13%, whereas YFV-17D-immune mice only lost approximately 3.5% of their body weight (Fig. 1B). By 4 dpi, the naïve group had lost approximately 14% of their body weight, significantly more than the immune group, which had already begun to recover after 2 dpi (Fig. 1B). Weight regain in YFV-17D-naïve mice was observed only after 4 dpi (Fig. 1B).

**Figure 1:**
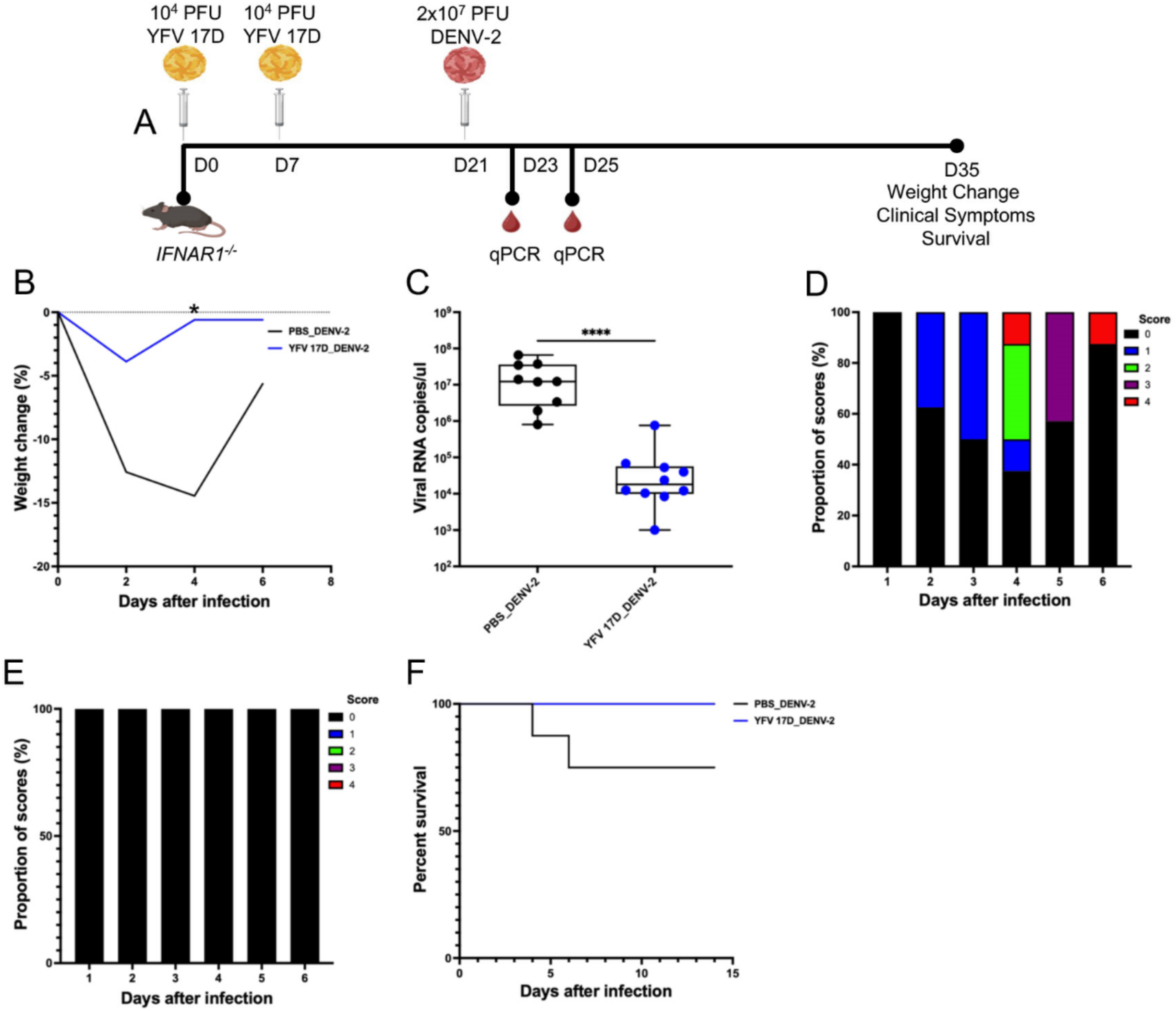
Heterologous immunity to YFV 17D provides protection against DENV-2. (A) Schematic diagram depicting YFV 17D immunization schedule followed by DENV-2 challenge. IFNAR1-/-mice were sham-immunized or immunized twice with YFV 17D, 7 days apart and subsequently challenged with 2×10^7^ PFU of DENV-2 at 14 days following the second immunization. Mice were monitored for viremia levels, weight change, symptoms and survival (B) Percentage change in body weight of YFV-naïve and YFV-immune mice following DENV-2 challenge until 6 days post-infection (dpi). Differences in weight loss over time were assessed using two-way ANOVA with a mixed-effect model for multiple comparisons. The data at each time point represents the mean weight of independent experiments (n=8 for YFV-naïve; n=7 for YFV-immune group). Significant difference was observed at 4dpi (P= 0.0331) (C) Levels of DENV-2 viremia in YFV-naïve and immune mice following DENV-2 challenge measured by quantitative RT-PCR at 2 dpi. Group differences were assessed by two-tailed Mann-Whitney test. Data are presented as mean ± SEM of independent experiments (n=9 for YFV-naïve; n=10 for YFV-immune group). Significant difference was observed at P<0.0001. Clinical symptoms of DENV-2 disease in (D) YFV-naïve (n=8) and (E) YFV-immune (n=7) mice observed until 6 dpi. Clinical scoring was based on a 5-point scale: 1, healthy; 2, ruffled fur; 3, ruffled fur and hunched posture; 4, ruffled fur, hunched posture and lethargy; 5, dead. (F) Survival outcomes of YFV-naïve (n=8) and YFV-immune (7) mice at 14 dpi. Differences between groups were assessed by the Gehan-Breslow-Wilcoxon test. P-values less than 0.05 were considered statistically significant. Non-significant differences were not indicated.

At 2 dpi, DENV-2 viremia levels in YFV-17D-naïve mice were significantly higher compared to YFV-17D-immune mice. (Fig. 1C). Viremia peaked at 2 dpi before declining, with some mice having undetectable viral loads by 4 dpi (Fig. 1C). While DENV-2 viremia levels decreased significantly from 2 dpi to 4 dpi in the naïve group, a similar but non-significant trend was observed in the immunized group (Fig. 1C). By 4 dpi, viremia levels were comparable between the two groups (Fig. 1C).

Clinically, YFV-17D-naïve mice exhibited progressive DENV-2-assocaited symptoms, worsening until 6 dpi, after which they began to recover (Fig. 1D, Supplementary Table 1). In contrast, all of the YFV-17D-immune mice remained asymptomatic throughout the infection (Fig. 1E). Finally, while all YFV-17D-immune mice survived, only 75% of naïve mice survived by 14 dpi, the endpoint of the study (Fig. 1F).

Collectively, these data demonstrate that pre-existing YFD-17D immunity significantly protects against DENV-2 infection, reducing weight loss, viral burden, and disease severity while improving survival outcomes.

### YFV 17D-specific antibodies do not neutralize nor enhance DENV-2 infection

Neutralizing antibodies are key correlates of vaccine-induced protection, including for the YFV-17D vaccine, which induces strong neutralizing responses against YFV re-infection (42). Therefore, to investigate whether YFV-17D-mediated cross-protection against DENV-2 is antibody-driven, we assessed the neutralizing and enhancing potential of YFV-17D-specific antibodies.

*IFNAR1^-/-^* mice were immunized twice with YFV-17D at a 7-day interval, and plasma was collected 14 days after the second immunization (Fig. 2A). As expected, YFV-17D-immune plasma antibodies exhibited strong neutralizing activity against YFV-17D (Fig. 2B). However, these antibodies failed to neutralize DENV-2, ZIKV, or WNV (Fig. 2B). Despite the lack of neutralization, the plasma antibodies from the YFV-17D-immune mice recognized and bound to the envelope (E) proteins of DENV-1, DENV-2, DENV-4, ZIKV, and WNV (Fig. 2C).

**Figure 2:**
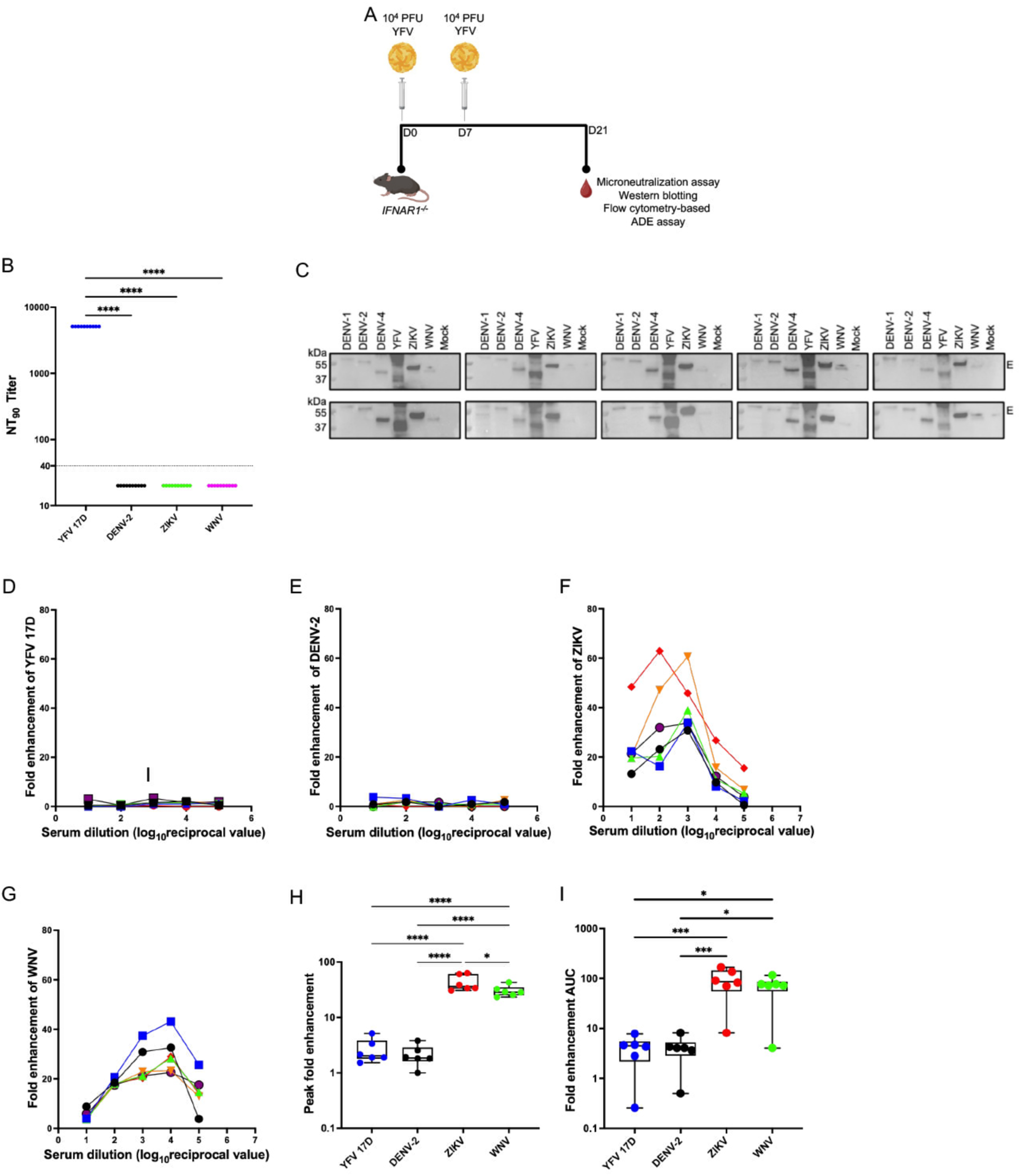
Heterologous antibodies to YFV neither neutralize nor enhance DENV-2 infection, but bind all orthoflaviviruses. (A) Schematic diagram depicting YFV-17D immunization schedule with serum collection performed 14 days after the second immunization. Sera from YFV-immune mice were used to determine neutralizing and antibody dependent enhancement (ADE) activities against YFV-17D, DENV-2, ZIKV and WNV, using a microneutralization assay and a flow cytometry-based ADE assay, respectively (B) Neutralizing antibody (nAb) titers against YFV-17D, DENV-2, ZIKV and WNV. Differences in antibody titers were assessed by the Kruskal-Wallis test with Dunn’s test for multiple comparisons. A significant difference was observed at P<0.0001. Data represent independent experiments (n=10). The dotted line shows the cut-off of neutralizing antibody titer. (C) Western blots demonstrating the binding of YFV-17D-specific antibodies to the envelope proteins of DENV-1, DENV-2, DENV-4, ZIKV and WNV. Cell lysates were prepared from virus-infected Vero cells, and viral protein components were resolved by SDS-PAGE. Blots were then probed with serum from YFV-17D-immune mice to assess antibody binding. The blots shown represent independent experiments n=10 YFV-17D immunized mice. Infection of U937 cells with (D) YFV 17D (E) DENV (F) ZIKV and (H) WNV in the presence of serially diluted serum from YFV-17D-immune mice or without serum. Data represent independent experiments (n=6) and are shown as the ratio of infected cells in the presence of serum to those without serum. E) Peak enhancement of infection of U937 cells with YFV-17D, DENV, ZIKV, and WNV using sera from YFV-17D-immune mice. Group differences were assessed using one-way ANOVA followed by Tukey’s multiple comparison test. Data are presented as mean ± SEM from independent experiments (n=6). Statistical significance was observed at P=0.0467 between YFV and ZIKV, and P<0.0001 for the other pairs. (I) Area under the curve (AUC), showing the total ADE activity of serum antibodies against YFV-17D, DENV-2, ZIKV and WNV. Group differences were assessed using one-way ANOVA followed by Holm-Šídák’s multiple comparisons test. Statistical significance was observed at P=0.01between WNV-YFV-17D and WNV-DENV-2 pairs, and P=0.001 for ZIKV-YFV-17D and ZIKV-DENV-2 pairs. Data are presented as mean ± SEM from independent experiments (n=6). P-values < 0.05 were considered statistically significant. Non-significant differences were not indicated.

Next, we assessed whether these non-neutralizing antibodies could mediate in vitro ADE. YFV-17D immune plasma antibodies did not enhance YFV-17D (Fig. 2D, 2H and 2I). Similarly, no enhancement of DENV-2 S221 was observed (Fig. 2E, 2H and 2I). Interestingly, however, YFV-17D plasma antibodies enhanced ZIKV (Fig. 2F, 2H and 2I) and WNV infections (Fig. 2G, 2H and 2I), highlighting differential in vitro ADE susceptibility among orthoflaviviruses.

Together, these findings indicate that YFV-17D-mediated cross-protection against DENV-2 is not antibody-driven, as YFV-17D-specific antibodies do not neutralize DENV-2 nor enhance its infection.

### YFV-17D-specific T cells cross-react with DENV antigens

To investigate whether T cells contribute to YFV-17D-mediated cross-protection, we assessed the cross-reactivity of YFV-17D-specific T cells against DENV antigens. Given the high homology between the orthoflavivirus structural and non-structural proteins (10), we hypothesized that YFV-17D-specific T cells might recognize DENV-derived epitopes. *IFNAR1^-/-^*mice were immunized twice with YFV-17D at a 7-day interval, and splenocytes were collected 14 days after the second immunization and pulsed with LFn-YFV or -DENV structural and non-structural proteins (Fig 3A). T cell responses were then evaluated via extracellular staining, followed by intracellular cytokine staining and flow cytometry, to quantify IFNγ-producing CD4^+^ and CD8^+^ T cells.

**Figure 3:**
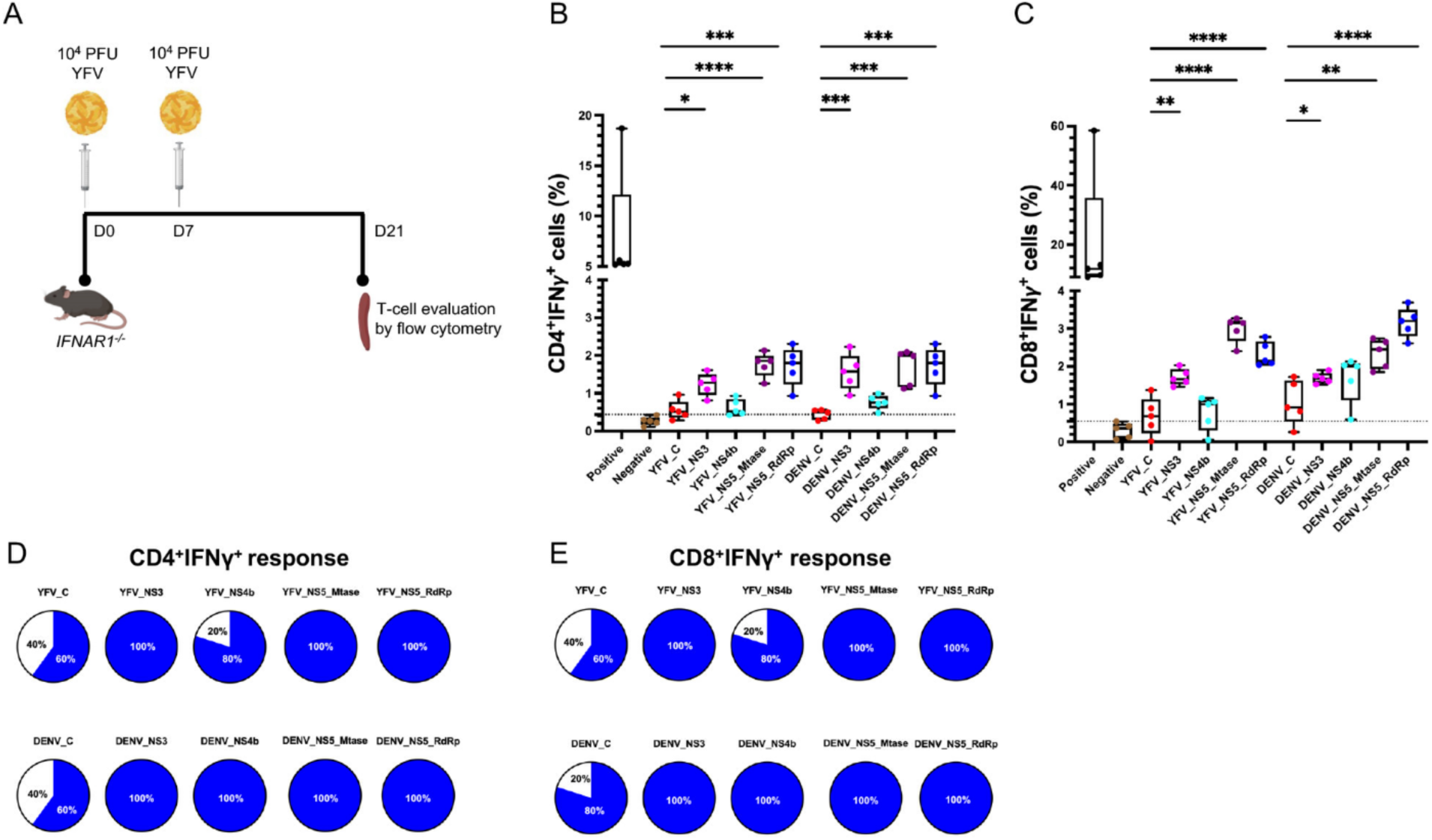
Cross-reactivity of YFV-specific T-cells to DENV. (A) Schematic diagram illustrating YFV-17D immunization schedule, with spleen collection performed 14 days after the second immunization. Splenocytes from YFV-17D-immune mice were analyzed by flow cytometry to assess cross-reactive YFV-specific T cell responses against structural (capsid) and non-structural (NS3, NS4b, NS5_RdRp, NS5_mTASE) proteins of DENV. (B) CD4^+^IFN-γ^+^ T cell responses to YFV and DENV structural and non-structural proteins. Data represents independent experiments (n=5). The dotted line represents the cutoff for CD4^+^IFN-γ^+^ T cell responses, determined by the receiver operating characteristic (ROC) curve analysis. The sensitivity and specificity values used to determine the cut-off value were 78% and 100%, respectively with a significant area under the curve (P=0.0033) (C) CD8^+^IFN-γ^+^ T cells response to YFV and DENV structural and non-structural proteins. The dotted line represents the cut-off for CD4^+^IFN-γ^+^ T cell responses, determined by the receiver operating characteristic (ROC) curve analysis. Data represents independent experiments (n=5). The sensitivity and specificity values used to determine the cut-off value were 78% and 100%, respectively with a significant area under the curve (P=0.0033) (D) Proportion of mice with positive CD4^+^IFN-γ^+^ T cell responses to the immunogens. Proportions were determined using the ROC-based curve cut-off value. The blue area of the pie charts represents proportion of mice with positive T cell responses, while the white area represents those with negative responses. (E) Proportion of mice with positive CD8^+^IFN-γ^+^ T cell responses to the immunogens. Proportions were determined based on the cut-off value generated from the ROC curve analysis. The blue area of the pie charts represents proportion of mice with positive T cell responses, while the white area represents those with negative responses.

As expected, and consistent with prior studies (9, 43), YFV-17D elicited robust CD4^+^IFNγ^+^ and CD8^+^IFNγ^+^ T cell responses against its non-structural proteins, NS3 and NS5, with significantly higher responses compared to capsid and NS4b (Fig. 3A-B). Notably, these CD4^+^IFNγ^+^ and CD8^+^IFNγ^+^ responses also extended to DENV-derived NS3 and NS5, showing significantly higher reactivity than against capsid and NS4b (Fig. 3A-B), indicating T cell cross-reactivity between these two orthoflaviviruses.

Using ROC curve analysis, we established cut-off values for CD4^+^IFNγ^+^ and CD8^+^IFNγ^+^ T cell responses (Supplementary Fig. 4), optimizing sensitivity and specificity for detecting responses to YFV and DENV proteins. 100% of mice exhibited CD4^+^IFNγ^+^ and CD8^+^IFNγ^+^ T cell response to NS3 and NS5 of both viruses (Fig. 3D-E), while responses to NS4b were 100% for DENV but 80% for YFV (Fig. 3D-E). Additionally, C responses ranged from 60-80% (Fig. 3D-E).

These findings suggest that orthoflavivirus infections elicit robust T cell responses primarily targeting conserved non-structural proteins, with YFV-17D-specific T cells also recognizing DENV antigens.

### YFV-17D-specific CD4^+^ and CD8^+^ T cells contribute to cross-protection against DENV-2

To determine whether YFV-17D-specific T cells mediate cross-protection, we depleted CD4^+^ and CD8^+^ T cells in YFV-immune mice at 3 and 1 days prior to DENV-2 S221 challenge, as well as 1 day post-challenge (Fig. 4A). Flow cytometric analysis confirmed nearly complete CD4^+^ (∼99%) and CD8^+^ (∼93%) T cell depletion (Fig. 4B-C).

**Figure 4:**
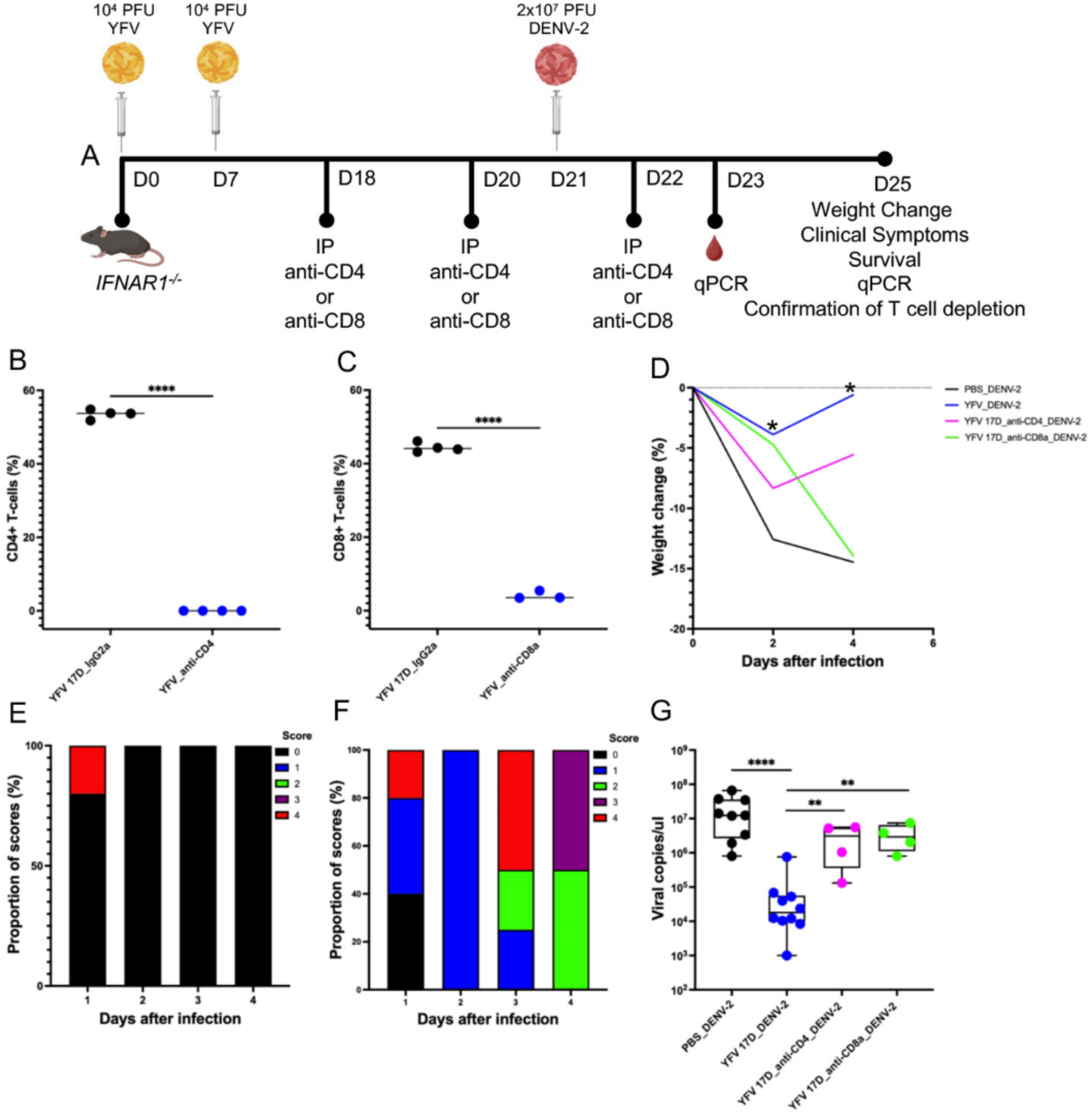
Depletion of CD4^+^ and CD8^+^ T cells abrogate YFV-17D-mediated cross-protection against DENV-2. (A) Schematic diagram depicting YFV-17D immunization and T cell depletion schedule, followed by DENV-2 challenge*. IFNAR1^-/-^* mice were sham-immunized or immunized twice with YFV-17D (7 days apart), followed by intraperitoneal injection of 200 µg of anti-CD4 or anti-CD8⍺ antibodies at 3 days and I day prior to challenge, as well as 1 day post-challenge with 2×10^7^ PFU of DENV-2. Mice were monitored for viremia levels, weight change, and clinical symptoms, and spleens were collected at 4 dpi to confirm CD4^+^ and CD8^+^T cell depletion. (B) CD4^+^ T cell depletion in YFV-17D-immune mice. (C) CD8^+^ T cell depletion in YFV-17D-immune mice. Data represents independent experiments (n=4). Group differences were assessed by the Mann-Whitney test, with statistical difference at P<0.0001. (D) Percentage change in body weight of YFV-17D-naïve, YFV-17D-immune, YFV-17D-immune with CD4^+^ or CD8^+^ T cell depletion post-challenge. Differences in weight loss over time were assessed using two-way ANOVA with a mixed-effect model for multiple comparisons. The data at each time point represents the mean weight of independent experiments (n=7 for YFV-17D-immune; n=8 for YFV-17D-naïve, n=4 each for n=4 each for YFV-17D-immune with CD4^+^ and CD8^+^ T cells depletion). Significant difference was observed at 2 dpi (P=0.0239) and 4 dpi (P=0.0258) between YFV-17D-immune with intact T cells and CD4 depletion group. Clinical symptoms of DENV-2 disease in YFV-17D-immune mice with (E) CD4 (n=4) and (F) CD8 (n=4) T cell depletion observed until 4 dpi. Clinical scoring was based on a 5-point scale: 1, healthy; 2, ruffled fur; 3, ruffled fur and hunched posture; 4, ruffled fur, hunched posture and lethargy; 5, dead. (G) Levels of DENV-2 viremia in YFV-17D-naïve, YFV-17D-immune, YFV-17D-immune with CD4^+^ or CD8^+^T cell depletion post-challenge, measured by quantitative RT-PCR at 2 dpi. Group differences were assessed by the Kruskal-Wallis test with Dunn’s test for multiple comparisons. Data are presented as mean ± SEM of independent experiments (n=10 for YFV-17D-immune; n=9 for YFV-17D-naïve, n=4 each for YFV-17D-immune with CD4^+^ and CD8^+^ T cells depletion). Significant difference was observed between YFV-17D-immune and naïve (P<0.0001), YFV-17D-immune with intact T cells and CD4^+^ depletion group (P=0.004), and YFV-17D-immune with intact T cells and CD8^+^ depletion group (P=0.002). P-values less than 0.05 were considered statistically significant. Non-significant differences were not indicated.

By 2 and 4 dpi, YFV-17D-immune mice with CD4^+^ T cell depletion experienced significantly less weight loss compared to those with intact T cell responses. In contrast, CD8^+^-depleted mice lost 14% of their body weight, comparable to YFV-17D-naïve mice, though the difference was not statistically significant (Fig. 4D).

Except for a mouse that succumbed at 1 dpi in CD4^+^-depleted group, the remaining mice exhibited no clinical signs of DENV-2 illness (Fig. 4E). In contrast, YFV-17D immune group with CD8^+^-depletion exhibited clinical symptoms associated with DENV-2 infection, which progressively worsened until 4 dpi, at which point the mice were euthanized (Fig. 4F, Supplementary Table 1). Moreover, at 2 dpi, DENV-2 viremia levels in both CD4^+^- or CD8^+^-depleted mice were significantly higher than in YFV-17D immune mice with intact T cell responses and were comparable to naïve mice (Fig 4G).

These findings demonstrate that YFV-17D-specific T cells play a critical role in cross-protection against DENV-2, with CD8^+^ T cells being particularly important in mitigating disease severity and viral burden.

### Cytotoxicity of YFV-17D-specific T-cells

Finally, to evaluate whether YFV-17D-specific T cells exhibit cytotoxicity against target cells pulsed with DENV-2 antigens, we performed an in vivo killing assay using CFSE-labeled splenocytes (Fig. 5A). Naïve splenocytes were pulsed with LFn-DENV NS3 or NS5 and subsequently stained with a high concentration of CFSE (CFSE^High^). These cells were then mixed at a 1:1 ratio with naïve splenocytes pulsed with LFn-LACV RdRp (irrelevant control immunogen) and stained with a low concentration of CFSE (CFSE^Low^). The mixed cells were then transferred into YFV-17D immune or naïve mice, and after 16 hours, in vivo killing of CFSE^High^ and CFSE^Low^ cells were measured by flow cytometry (Fig. 5A). The gating strategy is shown in Figure 5B.

**Figure 5:**
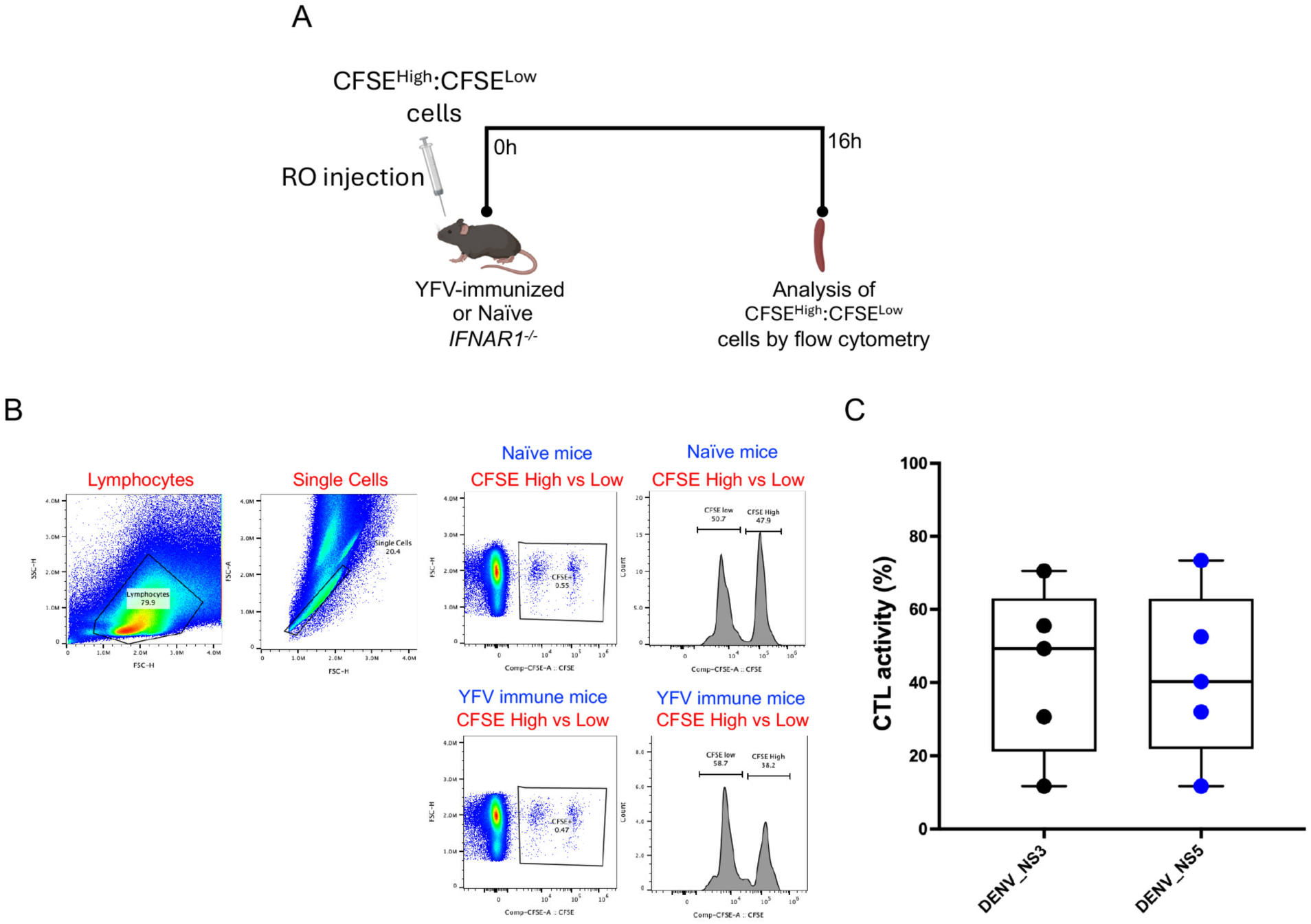
Cytotoxic activity of YFV-17D-specific T cells. (A) Schematic diagram illustrating adoptive transfer of a 1:1 mixture of CFSE^Low^ and CFSE^High^-pulsed splenocytes into YFV-17D-naïve or YFV-17D-immune mice. Splenocytes were isolated from YFV-17D-naïve and YFV-17D-immune mice. Naïve-derived cells were pulsed with LACV RdRp, while immune-derived cells were pulsed with either NS3 or NS5 proteins of DENV. 24 hrs later, NS3 and NS5-pulsed cells were stained with high concentration of CFSE (CFSE^High^), while LACV RdRp-pulsed cells stained with low concentration of CFSE (CFSE^Low^). CFSE^High^ and CFSE^Low^ cells were mixed in a 1:1 ratio and adoptive transferred by retro-orbital injection into YFV-17D naïve and immune mice. (B) Gating strategy for YFV-17D-specific T cell cytotoxic activity. Flow cytometry gating strategy used to identify splenocyte populations pulsed with either DENV NS3 or NS5 (CFSE^High^) and LACV RDRP (CFSE^Low^) in YFV-17D-immune and naïve mice. (C) In vivo cytotoxic activity of YFV-17D-specific T cells against splenocytes pulsed with NS3 and NS5 proteins of DENV. Data represents independent experiments (n=5). Group differences were assessed by two-tailed Mann-Whitney test. P-values less than 0.05 were considered statistically significant.

Strikingly, YFV-17D-specific T cells mediated ∼44% killing of DENV NS3-pulsed cells and ∼42% killing of NS5-pulsed cells (Fig. 5B). These findings demonstrate the cytotoxic activity of YFV-17D-specific T cells against cells pulsed with DENV antigens and provide functional evidence supporting T cell-mediated YFV-DENV cross-protection.

## Discussion

In this study, we demonstrate that pre-existing immunity to YFV-17D provides significant protection against DENV-2 S221 infection, as evidenced by reduced viral burden, weight loss, and disease severity, along with improved survival outcomes in mice. Investigation of the underlying mechanisms reveal that this protection is not mediated by cross-neutralizing antibodies against DENV-2. Instead, T cell depletion studies reveal that YFV-17D-specific CD4^+^ or CD8^+^ T cells play a pivotal role in cross-protection, as their depletion results in increased viral burden, weight loss, and disease severity. Furthermore, YFV-17D-specific T cells exhibit cytotoxic activity against cells pulsed with DENV non-structural antigens, confirming their role in YFV-17D-mediated cross-protection.

Currently, approximately 70 orthoflaviviruses have been identified into serocomplexes based on antigenic similarity and shared transmission vectors (44). Among these, nine including YFV, DENV-1-4, ZIKV, WNV, JEV, and tick-borne encephalitis virus (TBEV), are considered clinically relevant due to their rapid transmission, ability to cause severe disease or outbreaks, and significant contribution to the global disease burden (45). Given the widespread use of YFV-17D and JEV vaccines (45), combined with co-circulation of these orthoflaviviruses (34, 46) and increased global mobility, individuals in endemic areas face a heightened risk of multiple orthoflavivirus exposures. This has led to several epidemiological and animal studies investigating how heterologous immunity, acquired through either natural infection or vaccination, modulates immune responses and clinical outcomes upon subsequent exposure to a heterotypic orthoflavivirus (9, 11–24). The findings from these studies provide insight into the low transmission rates and varying clinical severity of certain orthoflaviviruses in specific regions, despite the presence of naïve populations and competent vectors, suggesting that pre-existing immunity may play a role in shaping orthoflavivirus epidemiology.

Notably, asymmetrical immunological interactions between DENV and ZIKV have been observed in epidemiological studies. Individuals with prior DENV immunity tend to experience milder or asymptomatic ZIKV infections (13), whereas those with pre-existing ZIKV immunity are at a higher risk of developing severe DENV disease (14, 17). Interestingly, DENV-immune serum or plasma has been shown to either protect or exacerbate ZIKV infections (11, 18, 26). In experimental studies using animal models, pre-existing ZIKV immunity was shown to protect or enhance dengue infections in mice (15, 20). Some studies have also demonstrated the protective effect of pre-existing DENV immunity against ZIKV infection in mice and rhesus macaques (12, 19, 21). Other studies have reported an increased ZIKV burden at the maternal-fetal interface in rhesus macaques (16), highlighting the complex and sometimes contradictory immune responses induced by these viruses.

Nonetheless, there is limited data on how pre-existing immunity to YFV modulates immune responses and clinical outcomes upon subsequent exposure to other clinically relevant orthoflaviviruses, and vice versa. For instance, a cohort study found no significant association between YFV-17D vaccination and severe dengue disease (7). Additionally, pre-existing YFV immunity in a murine model was shown to provide protection against subsequent ZIKV infections (8). Similarly, pre-existing immunity to DENV or ZIKV has been shown to provide protection against YFV in mice and macaques (23, 24). These findings have, to some extent, helped unravel the mystery surrounding the absence or low transmission rates of YFV in Asia and urban centers of South America, regions that are hyperendemic for DENV and ZIKV. This suggests that pre-existing DENV and/or ZIKV immunity may play a role in limiting YFV transmission in these regions.

Despite the substantial annual burden of orthoflavivirus infections in Sub-Saharan Africa, most outbreaks are mild or clinically inapparent (27). The introduction of the YFV-17D vaccine into routine immunization programs in this region has led to a significant proportion of the population developing immunity to the virus. These observations led us to the hypothesize that pre-existing immunity to YFV may contribute to limited occurrence of severe orthoflavivirus-related outbreaks in Sub-Saharan Africa. To investigate this, we employed a murine model to test whether pre-existing YFV-17D immunity confers cross-protection against DENV-2 infection.

Ultimately, we found that mice immune to YFV-17D were protected from secondary DENV-2 infection. YFV-17D immune mice exhibited significantly reduced viremia, disease severity, weight loss and improved survival outcomes compared to YFV-naïve mice. Despite neutralizing antibodies being the primary correlates of protection induced by the YFV 17D vaccine (42), plasma antibodies from YFV-17D-immune mice showed no neutralizing activity against DENV-2. This suggests that YFV-mediated cross-protection against DENV-2 is not driven by cross-neutralizing antibodies. Yet, the plasma antibodies exhibited cross-reactivity against DENV-2, prompting us to evaluate whether YFV non-neutralizing cross-reactive antibodies could exacerbate DENV-2 infection through ADE, a well-documented phenomenon in the context of sequential infections by DENV serocomplex viruses (47, 48), as well as JEV serocomplex viruses (49, 50). We found no evidence that these non-neutralizing antibodies enhanced DENV-2 infection in an in vitro assay. These findings are consistent with a similar study demonstrating YFV-mediated cross-protection against ZIKV (8) and JEV/YFV-17D chimeric virus (9), despite the absence of neutralizing antibodies, highlighting the potential role of T cell-mediated immunity.

Like heterologous antibody responses, orthoflavivirus cross-reactive T cell responses exhibit bidirectional immunomodulatory effects, either conferring protection or contributing to pathogenesis upon subsequent heterotypic infection, a phenomenon known as “original antigenic sin”. This occurs when cross-reactive T cells primed by the primary infecting virus drives an excessive cytokine response upon subsequent exposure to a heterotypic virus. Some studies have linked heterologous memory T cells to severe dengue disease (51). While the protective role of heterologous memory T cells has been extensively studied and well-characterized within the DENV serocomplex viruses (52–55), their impact, whether protective or pathogenic in the context of YFV and related-orthoflaviviruses, remains poorly understood.

To address this knowledge gap, we explored the potential role of T cells in YFV-mediated cross-protection against DENV-2 infections. We found that YFV-17D-specific T cells exhibit cross-reactivity to DENV, as evidenced by CD4^+^IFN^+^γ and CD8^+^IFNγ^+^ T cell responses to both structural and non-structural proteins of DENV. Notably, the response was stronger and more frequent for non-structural proteins, particularly NS3 and NS5, compared to the capsid protein. These findings suggest that YFV-17D-specific Τ cells may play a role in modulating immune responses and clinical outcomes against DENV infections.

To confirm this, we found that YFV-17D-immune mice depleted of CD4^+^ or CD8^+^ and subsequently infected with DENV-2 showed increased viral burden, weight loss and disease severity, compared to YFV-17D-immune mice with intact T cells. Furthermore, we demonstrated that YFV-17D-specific T cells exhibited cytotoxic activity against DENV, as evidenced by the killing of naïve splenocytes pulsed with DENV NS3 and NS5 proteins. Taken together, these data suggest that both YFV-17D-specific CD4^+^ and CD8^+^ T cells play a crucial role in mediating cross-protection against DENV-2 infection.

One of the strengths of this study is the identification of the immune component responsible for the protective effect of pre-existing YFV immunity against DENV-2 infection. Specifically, the identification of T cells as a key immune correlate of protection, despite the absence of cross-neutralizing antibodies, is consistent with findings from a similar study in which T cells from YFV-17D vaccinees were shown to reduce JEV/YFV-17D viremia to undetectable levels (9), even in the absence of neutralizing antibodies. Furthermore, the use of LFn-conjugated structural and non-structural proteins of YFV and DENV for cell stimulation enabled a more precise assessment of cellular responses against these viruses. Previous studies have relied on the use of in silico HLA-predicted peptides spanning small protein regions, which may exclude critical peptides required to elicit a robust T cell response (21, 22, 39, 56). In contrast, LFn facilitates the delivery of full-length proteins into the cytosol for native processing via the major histocompatibility complex, potentially eliciting a stronger T cell response compared to peptide-based approaches (33, 57, 58).

Despite these strengths, the study has several limitations. First, we utilized *IFNAR1^-/-^* mice, which may not fully recapitulate infection dynamics in immunocompetent humans or animals. Additionally, the use of an attenuated YFV strain may induce an immune response of a different magnitude compared to a wild-type strain, potentially altering the immune modulation of subsequent DENV infection. This may limit our understanding of how YFV immunity acquired through natural infection influences subsequent exposure to DENV. Furthermore, we only assessed the impact of YFV-17D immunity on DENV-2, without evaluating other DENV serotypes. Hence, the extent of immunological cross-reactivity between YFV and other serotypes remains uncertain, warranting further studies to elucidate these interactions. Indeed, a study found that YFV-immune mice were not able to provide cross-protection against DENV-1 challenge (59). This lack of protection, however, could be attributed to the single-dose immunization of C57B/6NTac mice, which likely is insufficient to confer immunity against DENV-1 infection. In contrast, the immunization schedule used in our study which provides significant protection against secondary DENV-2 infections, was adapted from a similar study demonstrating protection against ZIKV in mice with pre-existing YFV-17D immunity (8). This schedule also aligns with the dosage regimen previously administered to humans before 2015, when the advisory committee on immunization practice of CDC revised its recommendation to a single-dose vaccine (60). Our evaluation of YFV-17D immunity on subsequent DENV-2 infections in fourteen days following the second immunization may have also influenced the study outcomes. Future studies are needed to evaluate the long-term durability of YFV-17D immunity and its impact on DENV-2 infections. Moreover, the DENV-2 used is a lab-adapted strain which may also influence the findings, potentially limiting their generalizability to natural infections. Moreover, while we observed no cross-neutralizing antibodies against DENV-2, this does not rule out the protective potential role of non-neutralizing antibodies. These antibodies may exert effector functions such as antibody dependent cellular cytotoxicity or phagocytosis, which could contribute to controlling DENV-2 replication. Hence, further studies are needed to fully elucidate their role. Lastly, although CD4^+^- and CD8^+^-depleted YFV-immune mice showed increased levels of DENV-2 viremia, we did not delineate the specific contributions of CD4^+^ and CD8^+^ T-cells in controlling DENV-2 replication, an aspect previously explored in other orthoflavivirus sequential infection studies (52–55, 59). However, we demonstrated that YFV-17D-specific T cells exhibit cytotoxic activity against DENV-pulsed cells.

In conclusion, our study demonstrates that pre-existing immunity to YFV-17D provides significant protection against DENV-2 infection, despite the absence of cross-neutralizing antibodies. T cell depletion studies revealed that CD4^+^ and CD8^+^ T cells contribute to YFV-mediated cross-protection against DENV-2 infection. Furthermore, YFV-specific T cells exhibited cytotoxic activity against DENV-infected cells, confirming their protective role. These findings underscore the need for orthoflavivirus vaccine design that elicit robust T cell responses, which may confer cross-protection. Finally, these findings will necessitate further evaluation of heterologous immunity to orthoflavivirus and its role against subsequent heterotypic infections.

## Acknowledgements

We would like to thank Rutgers Global Health Institute, Rutgers Robert Wood Johnson Medical School, and The Child Health Institute of New Jersey for their continued support. This study received funding from New Jersey Health Foundation (IFH18-24 to BBH). This work was also supported by NIH NIAID grant (R01AI149502 to WKW).

## Author Contributions

PBT carried out most of the described experiments, analyzed experimental data, and wrote the original draft of manuscript. SG propagated and tittered the DENV-2 S221 and aided with conducting the in vitro ADE assays. RA aided with conducting the CFSE-based in vivo cytotoxicity assays. WKW prepared the orthoflavivirus Vero cell lysates.

BBH conceived the original idea, supervised the work, and planned and carried out experiments. All authors contributed to the interpretation of the data and played a role in reviewing and editing the manuscript.

## References

1. Pierson TC, Diamond MS. The continued threat of emerging flaviviruses. Nat Microbiol. 2020;5(6):796–812.

2. Sridhar S, Luedtke A, Langevin E, Zhu M, Bonaparte M, Machabert T, et al. Effect of Dengue Serostatus on Dengue Vaccine Safety and Efficacy. N Engl J Med. 2018;379(4):327–40.

3. Izmirly AM, Alturki SO, Alturki SO, Connors J, Haddad EK. Challenges in Dengue Vaccines Development: Pre-existing Infections and Cross-Reactivity. Front Immunol. 2020;11:1055.

4. Pattnaik A, Sahoo BR, Pattnaik AK. Current Status of Zika Virus Vaccines: Successes and Challenges. Vaccines. 2020;8(2):266.

5. Gubler DJ, Halstead SB. Is Dengvaxia a useful vaccine for dengue endemic areas? Bmj. 2019;367:l5710.

6. Thomas SJ, Yoon IK. A review of Dengvaxia®: development to deployment. Hum Vaccin Immunother. 2019;15(10):2295–314.

7. Luppe MJ, Verro AT, Barbosa AS, Nogueira ML, Undurraga EA, da Silva NS. Yellow fever (YF) vaccination does not increase dengue severity: A retrospective study based on 11,448 dengue notifications in a YF and dengue endemic region. Travel Med Infect Dis. 2019;30:25–31.

8. Vicente Santos AC, Guedes-da-Silva FH, Dumard CH, Ferreira VNS, da Costa IPS, Machado RA, et al. Yellow fever vaccine protects mice against Zika virus infection. PLoS Negl Trop Dis. 2021;15(11):e0009907.

9. Kalimuddin S, Tham CYL, Chan YFZ, Hang SK, Kunasegaran K, Chia A, et al. Vaccine-induced T cell responses control Orthoflavivirus challenge infection without neutralizing antibodies in humans. Nature Microbiology. 2025;10(2):374–87.

10. Rathore APS, St John AL. Cross-Reactive Immunity Among Flaviviruses. Front Immunol. 2020;11:334.

11. Dejnirattisai W, Supasa P, Wongwiwat W, Rouvinski A, Barba-Spaeth G, Duangchinda T, et al. Dengue virus sero-cross-reactivity drives antibody-dependent enhancement of infection with zika virus. Nature Immunology. 2016;17(9):1102–8.

12. Pantoja P, Pérez-Guzmán EX, Rodríguez IV, White LJ, González O, Serrano C, et al. Zika virus pathogenesis in rhesus macaques is unaffected by pre-existing immunity to dengue virus. Nat Commun. 2017;8:15674.

13. Rodriguez-Barraquer I, Costa F, Nascimento EJM, Nery NJ, Castanha PMS, Sacramento GA, et al. Impact of preexisting dengue immunity on Zika virus emergence in a dengue endemic region. Science. 2019;363(6427):607–10.

14. Katzelnick LC, Narvaez C, Arguello S, Lopez Mercado B, Collado D, Ampie O, et al. Zika virus infection enhances future risk of severe dengue disease. Science. 2020;369(6507):1123–8.

15. Shukla R, Shanmugam RK, Ramasamy V, Arora U, Batra G, Acklin JA, et al. Zika virus envelope nanoparticle antibodies protect mice without risk of disease enhancement. EBioMedicine. 2020;54:102738.

16. Crooks CM, Weiler AM, Rybarczyk SL, Bliss MI, Jaeger AS, Murphy ME, et al. Previous exposure to dengue virus is associated with increased Zika virus burden at the maternal-fetal interface in rhesus macaques. PLoS Negl Trop Dis. 2021;15(7):e0009641.

17. Estofolete CF, Versiani AF, Dourado FS, Milhim B, Pacca CC, Silva GCD, et al. Influence of previous Zika virus infection on acute dengue episode. PLoS Negl Trop Dis. 2023;17(11):e0011710.

18. Tan JY, Tan KK, AbuBakar S. DIFFERENTIAL CROSS-NEUTRALIZATION OF DENGUE IMMUNE SERA AGAINST ZIKA VIRUS. International Journal of Infectious Diseases. 2023;134:S8.

19. Wen J, Elong Ngono A, Regla-Nava JA, Kim K, Gorman MJ, Diamond MS, et al. Dengue virus-reactive CD8(+) T cells mediate cross-protection against subsequent Zika virus challenge. Nat Commun. 2017;8(1):1459.

20. Fowler AM, Tang WW, Young MP, Mamidi A, Viramontes KM, McCauley MD, et al. Maternally Acquired Zika Antibodies Enhance Dengue Disease Severity in Mice. Cell Host Microbe. 2018;24(5):743–50.e5.

21. Regla-Nava JA, Elong Ngono A, Viramontes KM, Huynh AT, Wang YT, Nguyen AT, et al. Cross-reactive Dengue virus-specific CD8(+) T cells protect against Zika virus during pregnancy. Nat Commun. 2018;9(1):3042.

22. Grifoni A, Pham J, Sidney J, O’Rourke PH, Paul S, Peters B, et al. Prior Dengue Virus Exposure Shapes T Cell Immunity to Zika Virus in Humans. J Virol. 2017;91(24).

23. Shinde DP, Plante JA, Scharton D, Mitchell B, Walker J, Azar SR, et al. Potential role of heterologous flavivirus immunity in preventing urban transmission of yellow fever virus. Nature Communications. 2024;15(1):9728.

24. Shinde DP, Walker J, Reyna RA, Scharton D, Mitchell B, Dulaney E, et al. Mechanisms of Flavivirus Cross-Protection against Yellow Fever in a Mouse Model. Viruses. 2024;16(6).

25. Agrawal B. Heterologous Immunity: Role in Natural and Vaccine-Induced Resistance to Infections. Front Immunol. 2019;10:2631.

26. Bardina SV, Bunduc P, Tripathi S, Duehr J, Frere JJ, Brown JA, et al. Enhancement of Zika virus pathogenesis by preexisting antiflavivirus immunity. Science. 2017;356(6334):175–80.

27. Khan A, Ndenga B, Mutuku F, Bosire CM, Okuta V, Ronga CO, et al. Majority of pediatric dengue virus infections in Kenya do not meet 2009 WHO criteria for dengue diagnosis. PLOS Glob Public Health. 2022;2(4):e0000175.

28. Liang Y, Dai X. The global incidence and trends of three common flavivirus infections (Dengue, yellow fever, and Zika) from 2011 to 2021. Front Microbiol. 2024;15:1458166.

29. WHO. Disease Outbreak News; Dengue-Global situation 2024 [Available from: https://www.who.int/emergencies/disease-outbreak-news/item/2023-DON518.

30. Bhatt S, Gething PW, Brady OJ, Messina JP, Farlow AW, Moyes CL, et al. The global distribution and burden of dengue. Nature. 2013;496(7446):504-7.

31. Kuno G. The Absence of Yellow Fever in Asia: History, Hypotheses, Vector Dispersal, Possibility of YF in Asia, and Other Enigmas. Viruses. 2020;12(12).

32. Herrera BB, Chang CA, Hamel DJ, Mboup S, Ndiaye D, Imade G, et al. Continued Transmission of Zika Virus in Humans in West Africa, 1992-2016. J Infect Dis. 2017;215(10):1546–50.

33. Herrera BB, Tsai WY, Chang CA, Hamel DJ, Wang WK, Lu Y, et al. Sustained Specific and Cross-Reactive T Cell Responses to Zika and Dengue Virus NS3 in West Africa. J Virol. 2018;92(7).

34. Gallon S, Sy M, Tonto PB, Ndiaye IM, Toure M, Gaye A, et al. Seroprevalence of dengue, Zika, yellow fever and West Nile viruses in Senegal, West Africa. medRxiv. 2024:2024.11.21.24317738.

35. Lei C, Yang J, Hu J, Sun X. On the Calculation of TCID(50) for Quantitation of Virus Infectivity. Virol Sin. 2021;36(1):141–4.

36. Alm E, Lindegren G, Falk KI, Lagerqvist N. One-step real-time RT-PCR assays for serotyping dengue virus in clinical samples. BMC Infectious Diseases. 2015;15(1):493.

37. Tonto PB, Sy M, Ndiaye IM, Toure M, Gaye A, Aidara M, et al. Seroprevalence of Chikungunya and O’nyong-nyong Viruses in Senegal, West Africa. J Med Virol. 2025;97(3):e70282.

38. Laposova K, Oveckova I, Tomaskova J. A simple method for isolation of cell-associated viral particles from cell culture. J Virol Methods. 2017;249:194–6.

39. Elong Ngono A, Young MP, Bunz M, Xu Z, Hattakam S, Vizcarra E, et al. CD4+ T cells promote humoral immunity and viral control during Zika virus infection. PLoS Pathog. 2019;15(1):e1007474.

40. Habibzadeh F. Data Distribution: Normal or Abnormal? J Korean Med Sci. 2024;39(3):e35.

41. Pan P, Du X, Zhou Q, Cui Y, Deng X, Liu C, et al. Characteristics of lymphocyte subsets and cytokine profiles of patients with COVID-19. Virol J. 2022;19(1):57.

42. Wieten RW, Jonker EF, van Leeuwen EM, Remmerswaal EB, Ten Berge IJ, de Visser AW, et al. A Single 17D Yellow Fever Vaccination Provides Lifelong Immunity; Characterization of Yellow-Fever-Specific Neutralizing Antibody and T-Cell Responses after Vaccination. PLoS One. 2016;11(3):e0149871.

43. James EA, LaFond RE, Gates TJ, Mai DT, Malhotra U, Kwok WW. Yellow fever vaccination elicits broad functional CD4+ T cell responses that recognize structural and nonstructural proteins. J Virol. 2013;87(23):12794–804.

44. Calisher CH, Karabatsos N, Dalrymple JM, Shope RE, Porterfield JS, Westaway EG, et al. Antigenic relationships between flaviviruses as determined by cross-neutralization tests with polyclonal antisera. J Gen Virol. 1989;70 (Pt 1):37–43.

45. Heinz FX, Stiasny K. Flaviviruses and flavivirus vaccines. Vaccine. 2012;30(29):4301–6.

46. Ferenczi E, Bán E, Ábrahám A, Kaposi T, Petrányi G, Berencsi G, et al. Severe tick-borne encephalitis in a patient previously infected by West Nile virus. Scandinavian Journal of Infectious Diseases. 2008;40(9):759–61.

47. Morens DM, Halstead SB. Measurement of antibody-dependent infection enhancement of four dengue virus serotypes by monoclonal and polyclonal antibodies. J Gen Virol. 1990;71 (Pt 12):2909–14.

48. Vaughn DW, Green S, Kalayanarooj S, Innis BL, Nimmannitya S, Suntayakorn S, et al. Dengue viremia titer, antibody response pattern, and virus serotype correlate with disease severity. J Infect Dis. 2000;181(1):2–9.

49. Lobigs M, Larena M, Alsharifi M, Lee E, Pavy M. Live chimeric and inactivated Japanese encephalitis virus vaccines differ in their cross-protective values against Murray Valley encephalitis virus. J Virol. 2009;83(6):2436–45.

50. Lobigs M, Pavy M, Hall R. Cross-protective and infection-enhancing immunity in mice vaccinated against flaviviruses belonging to the Japanese encephalitis virus serocomplex. Vaccine. 2003;21(15):1572–9.

51. Mongkolsapaya J, Dejnirattisai W, Xu XN, Vasanawathana S, Tangthawornchaikul N, Chairunsri A, et al. Original antigenic sin and apoptosis in the pathogenesis of dengue hemorrhagic fever. Nat Med. 2003;9(7):921–7.

52. Elong Ngono A, Chen HW, Tang WW, Joo Y, King K, Weiskopf D, et al. Protective Role of Cross-Reactive CD8 T Cells Against Dengue Virus Infection. EBioMedicine. 2016;13:284–93.

53. Weiskopf D, Angelo MA, de Azeredo EL, Sidney J, Greenbaum JA, Fernando AN, et al. Comprehensive analysis of dengue virus-specific responses supports an HLA-linked protective role for CD8+ T cells. Proc Natl Acad Sci U S A. 2013;110(22):E2046–53.

54. Yauch LE, Prestwood TR, May MM, Morar MM, Zellweger RM, Peters B, et al. CD4+ T cells are not required for the induction of dengue virus-specific CD8+ T cell or antibody responses but contribute to protection after vaccination. J Immunol. 2010;185(9):5405–16.

55. Yauch LE, Zellweger RM, Kotturi MF, Qutubuddin A, Sidney J, Peters B, et al. A protective role for dengue virus-specific CD8+ T cells. J Immunol. 2009;182(8):4865–73.

56. Grifoni A, Voic H, Dhanda SK, Kidd CK, Brien JD, Buus S, et al. T Cell Responses Induced by Attenuated Flavivirus Vaccination Are Specific and Show Limited Cross-Reactivity with Other Flavivirus Species. J Virol. 2020;94(10).

57. Herrera BB, Hamel DJ, Oshun P, Akinsola R, Akanmu AS, Chang CA, et al. A modified anthrax toxin-based enzyme-linked immunospot assay reveals robust T cell responses in symptomatic and asymptomatic Ebola virus exposed individuals. PLoS Negl Trop Dis. 2018;12(5):e0006530.

58. Herrera BB, Chaplin B, S MB, Abdullahi A, He M, Fisher SM, et al. Pre-pandemic cross-reactive antibody and cellular responses against SARS-CoV-2 among female sex workers in Dakar, Senegal. Front Public Health. 2025;13:1522733.

59. Saron WAA, Rathore APS, Ting L, Ooi EE, Low J, Abraham SN, et al. Flavivirus serocomplex cross-reactive immunity is protective by activating heterologous memory CD4 T cells. Sci Adv. 2018;4(7):eaar4297.

60. CDC. Yellow Fever Vaccine Booster Doses: Recommendations of the Advisory Committee on Immunization Practices 2015 [Available from: https://www.cdc.gov/mmwr/preview/mmwrhtml/mm6423a5.htm.

