## Supplementary Material for "Pre-existing YFV-17D immunity mediates T cell cross-protection against DENV-2 infection"

Supplementary Table 1: Clinical scoring of DENV-2 disease

| Clinical score | Symptoms |
|---|---|
| 1 | Healthy |
| 2 | Ruffled fur |
| 3 | Ruffled fur and hunched posture |
| 4 | Ruffled fur, hunched posture and lethargy |
| 5 | Death |

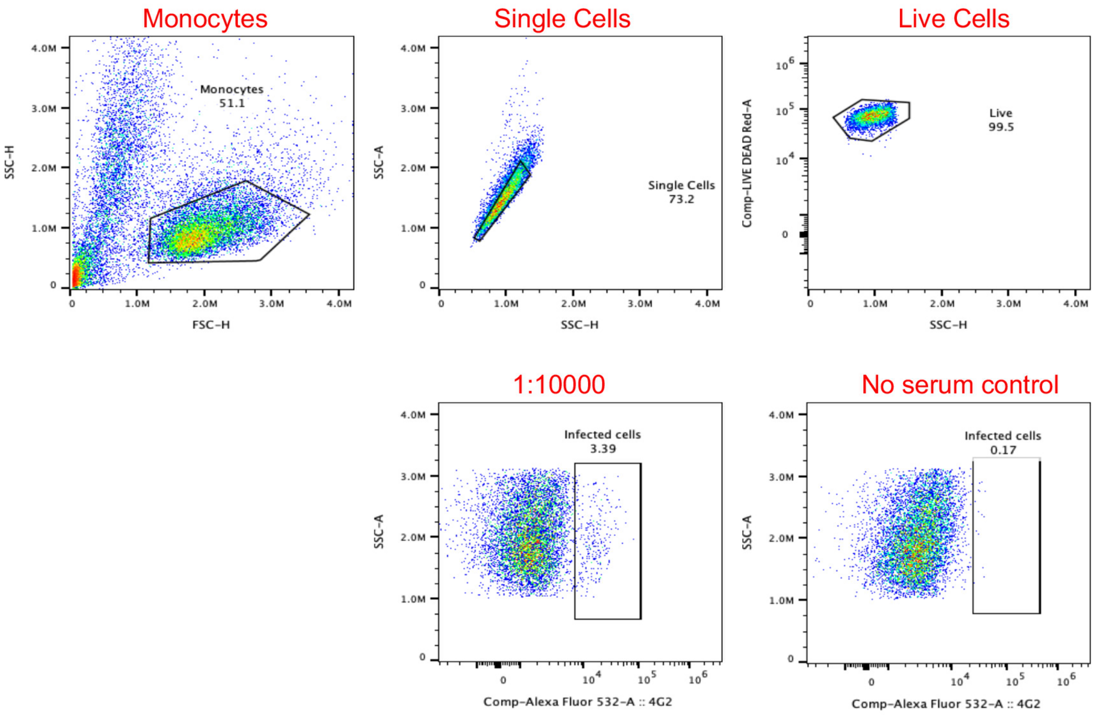


Supplementary Figure 1: Gating strategy for the in vitro ADE activity. Flow cytometry gating strategy used to identify U937 cell population infected with YFV-17D, DENV-2, ZIKV and WNV.

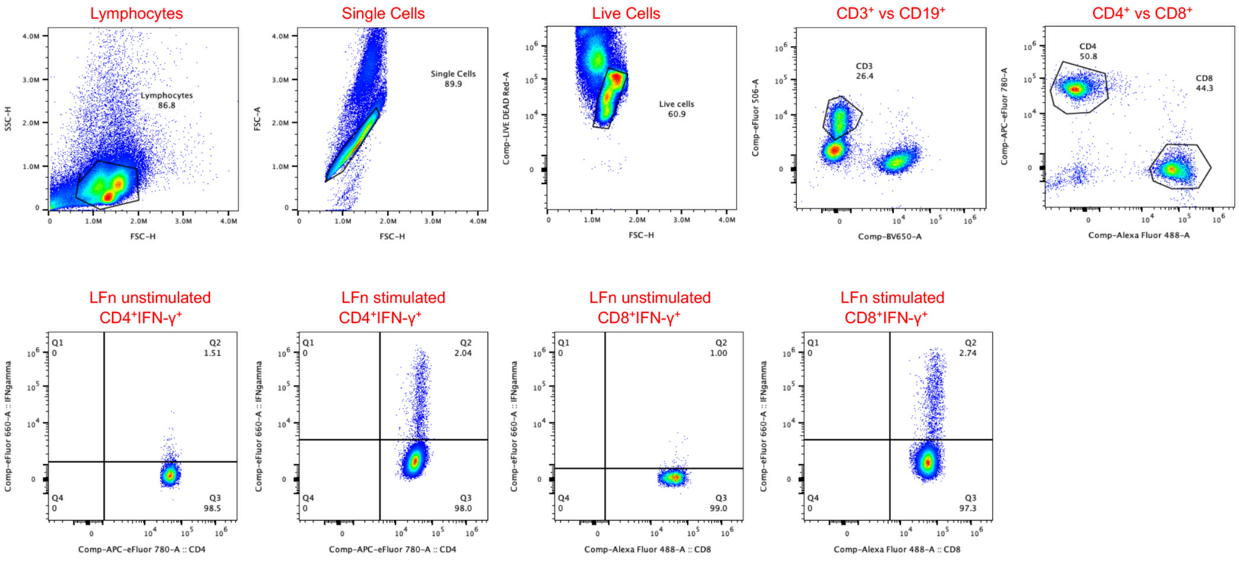


Supplementary Figure 2: Gating strategy for YFV-17D-specific T cells cross-reactivity to DENV. Flow cytometry gating strategy used to identify CD4$^{+}$IFN$\gamma^{+}$ and CD8$^{+}$IFN$\gamma^{+}$ T cell populations responding to structural (capsid) and non-structural (NS3, NS4b, NS5_RdRp, NS5_mTASE) proteins of YFV and DENV.

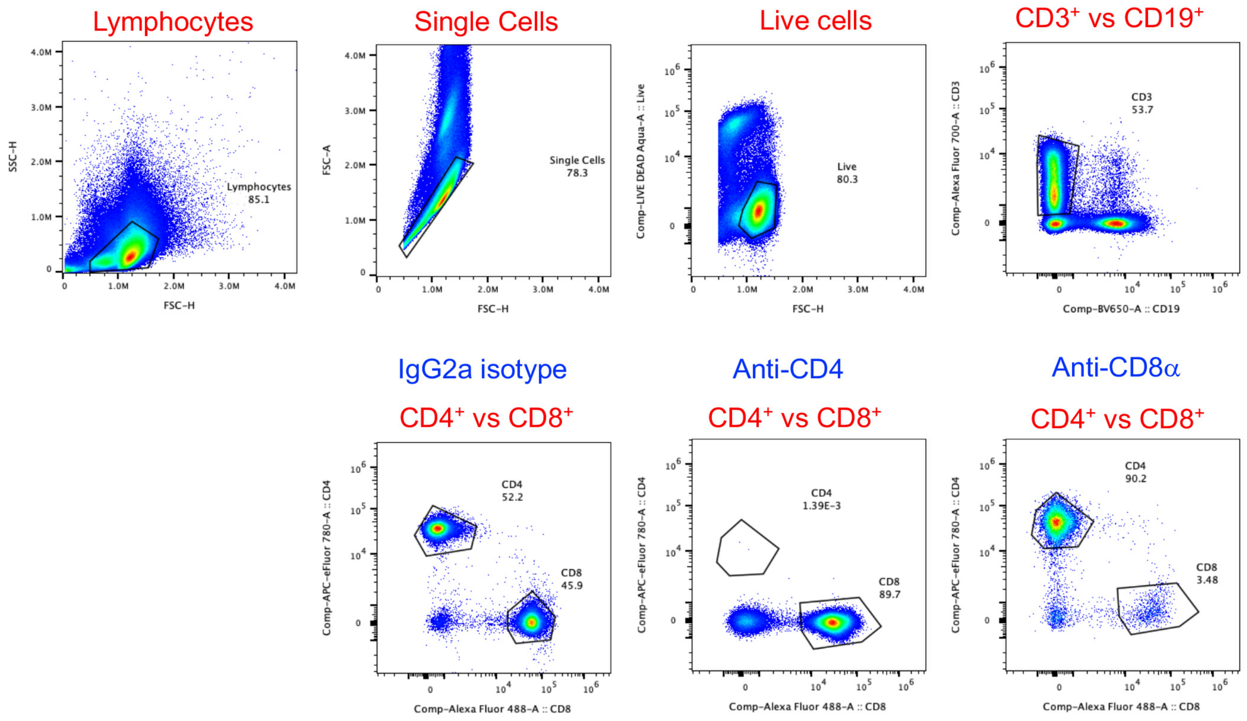


Supplementary Figure 3: Gating strategy for confirming $CD4^{+}$ and $CD8^{+}$ T cells depletion in YFV-17D-immune mice. Flow cytometry gating strategy used to assess the efficiency of $CD4^{+}$ and $CD8^{+}$ T cell depletion in mice following YFV-17D immunization

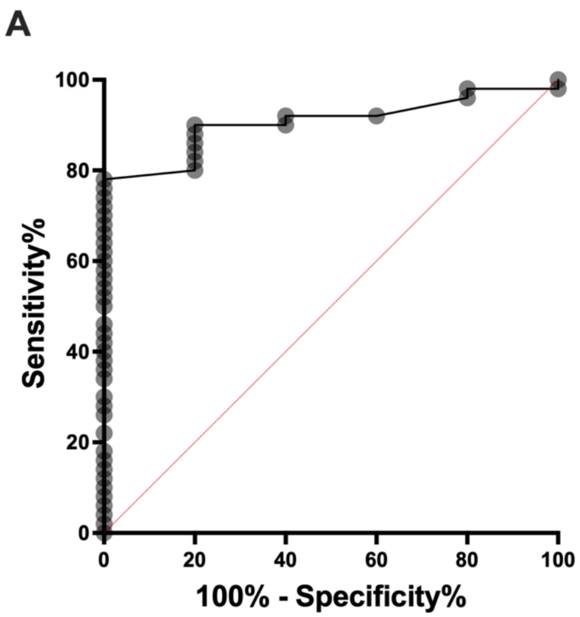


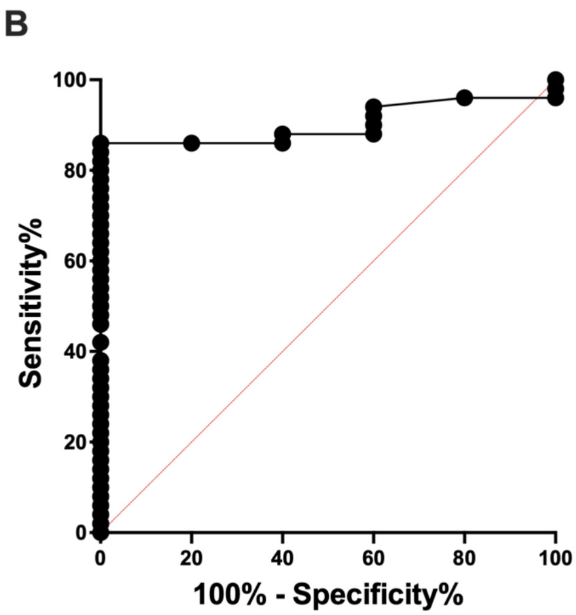


| | |
|---|---|
| AUC (95% CI) | 0.902 (0.8194-0.9846) |
| p value | 0.0033 |
| Sensitivity (95% CI) | 78 (64.76-87.25) |
| Specificity (95% CI) | 100 (56.55-100) |
| Cut-off | 0.435 |

| | |
|---|---|
| AUC (95% CI) | 0.902 (0.8194-0.9846) |
| p value | 0.0033 |
| Sensitivity (95% CI) | 86 (73.81-93.05) |
| Specificity (95% CI) | 100 (56.55-100) |
| Cut-off | 0.545 |

Supplementary Figure 4: Determination of T-cell response. (A) Receiver operating characteristic (ROC) curve analysis used to determine the threshold value for positive $CD4^{+}IFN\gamma^{+}$ T-cell response to structural (capsid) and non-structural (NS3, NS4b, NS5_RdRp, NS5_mTASE) proteins of YFV and DENV. (B) ROC curve analysis used to determine the threshold value for positive $CD8^{+}IFN\gamma^{+}$ T-cell response to structural (capsid) and non-structural (NS3, NS4b, NS5_mTASE, NS5_RdRp) proteins of YFV and DENV.
